# Unraveling the Enzymatic Mechanism of the SARS-CoV-2 RNA-Dependent-RNA-Polymerase. An Unusual Active Site Leading to High Replication Rates

**DOI:** 10.1101/2022.02.02.478873

**Authors:** Emmanuelle Bignon, Antonio Monari

## Abstract

Viral infection relies on the hijacking of cellular machineries to enforce the reproduction of the infecting virus and its subsequent diffusion. In this context the replication of the viral genome is a key step performed by specific enzymes, i.e. polymerases. The replication of SARS-CoV-2, the causative agent of the COVID-19 pandemics, is based on the duplication of its RNA genome, an action performed by the viral RNA-dependent-RNA polymerase. In this contribution, for the first time and by using two-dimensional enhanced sampling quantum mechanics/ molecular mechanics, we have determined the chemical mechanisms leading to the inclusion of a nucleotide in the nascent viral RNA strand. We prove the high efficiency of the polymerase, which lowers the activation free energy to less than 10 kcal/mol. Furthermore, the SARS-CoV-2 polymerase active site is slightly different from those found usually found in other similar enzymes, and particularly it lacks the possibility to enforce a proton shuttle via a nearby histidine. Our simulations show that this absence is partially compensate by lysine, whose proton assist the reaction opening up an alternative, but highly efficient, reactive channel. Our results present the first mechanistic resolution of SARS-CoV-2 genome replication and shed light on unusual enzymatic reactivity paving the way for future rational design of antivirals targeting emerging RNA viruses.

## Introduction

The emergence of a new *β*−coronavirus, observed for the first time at the end of 2019 in Wuhan, China, has led to the sudden outbreak of the COVID-19 pandemic.^1–5^ By the beginning of 2022, COVID-19 has affected almost all the continents, and has pushed public governments to implement severe containment and social distancing measures, seriously affecting economic and social life. The novel *β*−coronavirus has been recognized as the causative agent of a severe acute respiratory stress (SARS) and thus named SARS-CoV-2, to point out its similarity with SARS-CoV whose outbreak dated back to 2003. SARS-CoV-2 also shares structural and phylogenetic analogy with the Middle East Respiratory Syndrome Coronavirus (MERS-CoV) emerged in 2012. Differently from its predecessors, SARS-CoV-2 has a lower mortality ratio (estimated at around 2-3 %)^6–8^ and in some cases it develops asymptomatically, especially for its latest B.1.1.519 (omicron) variant.^9–11^ However, its high contagiousness and the possible development of severe outcomes, especially in elder patients or subjects presenting comorbidities, has caused and still causes an impressive stress on the health systems, only mitigated by the development of novel vaccines also including the messenger RNA (mRNA) strategy.^12–14^

As all *β*−coronaviruses, SARS-CoV-2 is an enveloped, rather large, positive-sense RNA virus.^15^ As such its genetic material is contained inside a virion protected by a lipid membrane, and is constituted by a positive 5’ to 3’ single-stranded RNA consisting of about 30,000 nucleobases.^16^ Upon infection the viral genetic material is injected into the cells where it hijacks the cellular machinery to enforce the production of viral proteins before initiating its replication. The analysis of SARS-CoV-2 transcriptome^17^ shows that in addition to structural proteins, including the highly celebrated Spike protein,^18^ which form the viral envelope and allow the interactions with human receptors,^19,20^ two large polyproteins are expressed and self-cleaved by the two viral enzymes, i.e. the 3CL-like (3CL^*PRO*^) and the papaine-like (PL^*PRO*^) proteases.^21–23^ This leads to the maturation of several non-structural proteins (NSP) which exert different (enzymatic) functions allowing SARS-CoV-2 to complete its maturation and assembly into novel virions, while also eluding the innate cellular immune system. Among those the RNA-dependent-RNA-polymerase (RNAP), composed of the complex NSP12/NSP7, is of particular importance.^24,25^ It uses the original viral positive-sense RNA strand to produce an intermediate negative-sense strand which is further used as a template allowing the synthesis of a novel duplicated positive-sense RNA strand. Of note, the RNA replication takes place in the cytoplasm in double-membrane vesicles and differently from other RNA viruses, SARS-CoV-2 also disposes of a replication proof-reading system based on a viral exonuclease,^26,27^ which limits replication errors and considerably slows down its mutation rate. Obviously SARS-CoV-2 RNAP constitutes an ideal target for possible pharmacological development.,^28–30^ Its inhibition would, indeed, result in the arrest of the viral replication cycle and of the infection. As a matter of fact, some nucleotide analogous,^29,30^ also used against some retroviruses such as Human Immunodeficiency Virus (HIV), have been proposed for SARS-CoV-2 treatment and while they have shown interesting *in vitro* and *in vivo* efficiency their clinical use is strongly limited also by rather sever side-effects and a non-trivial pharmacokinetic and administration. Between the proposed nucleotide analogous inhibiting SARS-CoV-2, we may cite Remdesivir,^31^ acting by impeding the RNA translocation,^32^ which follows the inclusion of a new nucleotide to free the active site and slide along the template, and Favipiravir,^33,34^ which instead directly inhibits the RNAP catalytic site.^35^ While the structure of SARS-CoV-2 RNAP has been resolved,^24,35^ also in presence of inhibitors, the studies focusing on the enzymatic mechanism from a molecular and biochemical point of view are rare. Yet, the precise characterization of the rate limiting steps of the enzymatic process, together with the identification of the structures of the key intermediates and transition states would be of invaluable help in the design of efficient inhibitors.

From a chemical standpoint, polymerases are metalloenzymes, sharing many structural and chemical similarities with exo- and endonucleases. Their active site is usually constituted by a bimetallic 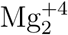 cluster which is stabilized by the interaction with hard electron donors such as the anionic oxygen atoms of aspartates (Figure 1). Furthermore, the presence of histidine residues which in addition to enhance the stabilization of the bimetallic cluster may also participate in proton shuttle completing the catalytic cycle, is also widely conserved and appears as fundamental.^36^ Furthermore, and upon reactive conformations, the 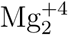 cluster is also interacting with the 3’ oxygen of the nascent nucleic acid strand (3’) and with the triphosphate form of the incorporating nucleotide. The selectivity and replication accuracy are assured by the fact that the triphosphate reactant nucleotide is Watson and Crick paired with the complementary base on the template strand. It is generally accepted that the 3’ OH group of the nascent strand is easily deprotonated, also thanks to its activation provided by 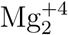, either by close-by acidic residues or by environmental water or OH^*−*^. The incorporation of the nucleotide is obviously accompanied by the cleavage of a P-O bonds on the triphosphate reactant. The release of the pyrophosphate intermediate and its protonation, usually assured by a catalytic histidine, are also necessary to complete the cycle.^36^ If SARS-CoV-2 RNAP shares strong similarities with other polymerases, particularly the presence of the 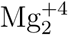 cluster, its active site is nonetheless peculiar due to the absence of histidines in close proximity of the enzymatic pocket. This suggests that the precise chemical mechanisms leading to the inclusion of the triphosphate nucleotide in the nascent RNA strand may be different from those commonly admitted in analogous systems. In this contribution, starting from the structure of the SARS-CoV-2 RNAP in complex with a RNA double strand resolved by Naydenova et al.^35^ (pdb code 7AAP, see Figure 1-A) we apply molecular modeling and simulations technique using both classical and hybrid quantum mechanical / molecular mechanical (QM/MM) strategies to unravel both the structural features and the chemical mechanism of RNAP, which is crucial for the replication of SARS-CoV-2. More specifically, by using two-dimensional (2D) enhanced sampling at QM/MM level we reveal from the one-side the low activation barrier of the polymerase-catalyzed reaction, as well as the fundamental role of a lysine assisting the reaction and efficiently compensating for the absence of catalytic histidines nearby the active site.

**Figure 1:**
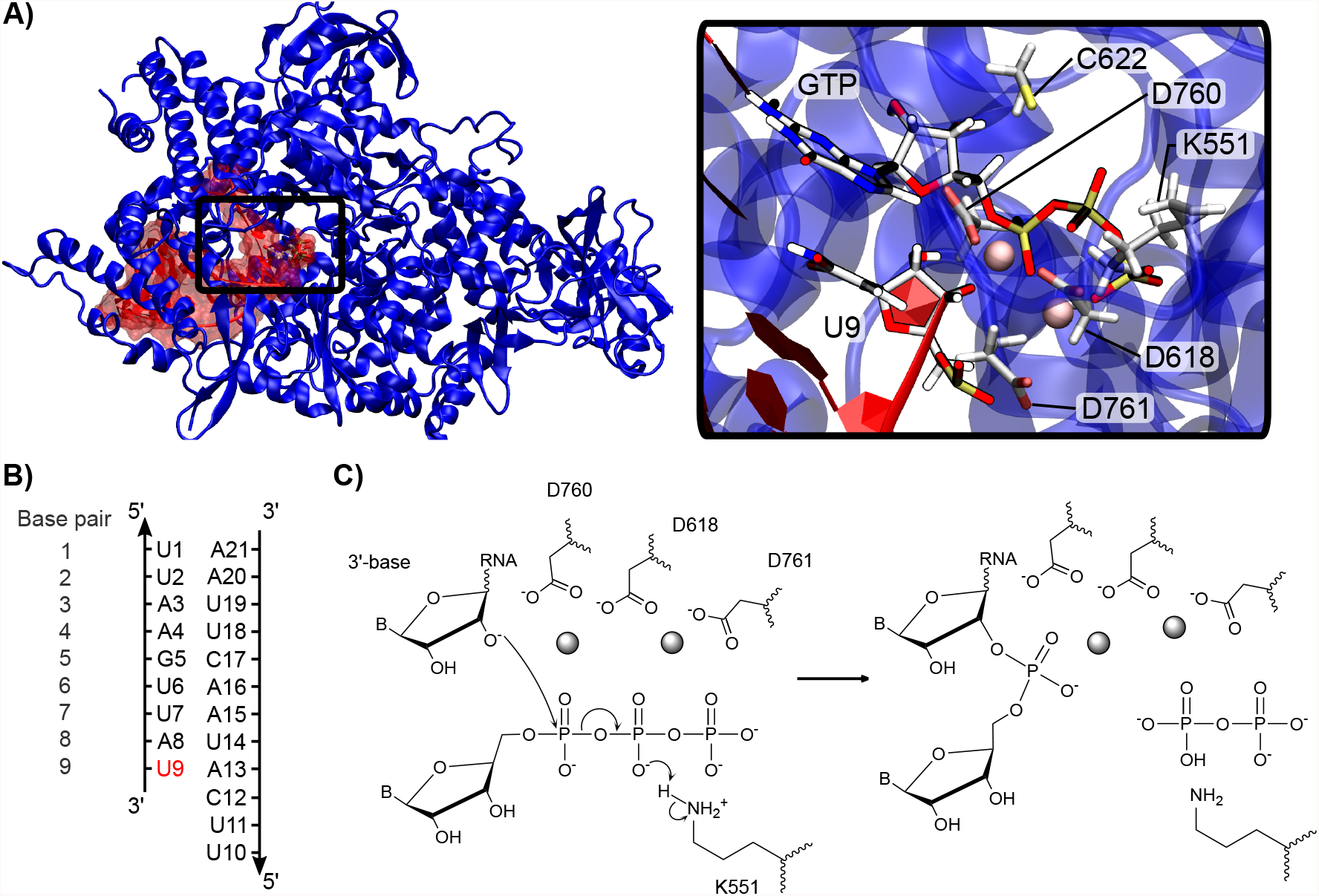
A) Representative structure extracted from the classical MD simulation of the RNAP (blue) interacting with a RNA double-strand (red) composed of the template and the nascent strand. A zoom of the active site is also provided highlighting GTP, the 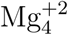 cluster, and key protein residues such as K551. B) Sequence of the RNA strand used consistently for all the simulation. C) proposed chemical mechanisms for the inclusion of GTP on the nascent RNA strand.

## Methods

All classical molecular dynamics (MD) simulations were carried on using NAMD.^37,38^ QM/MM simulations have been performed using the Terachem code^39^ interfaced with Amber 16.^40^ Visualization, rendering of MD trajectories and plots were performed using the VMD,^41^ Gnuplot and RStudio softwares.

### System setup and classical MD simulations

The starting structure was generated from the crystal structure of SARS-CoV-2 RNA-dependent RNA polymerase complexed with the Favipiravir inhibitor (PDB ID 7AAP). The Favipiravir moiety was *in silico* mutated to the reactive Guanine Triphosphate (GTP), keeping the catalytic magnesium ions as positioned in the crystal structure. As we chose to focus on the reaction step involving the attack of the deprotonated 3’ terminal nucleotide (rU9) of the synthesized RNA strand, the O3’ was coherently kept deprotonated. Force field parameters of the 3’ deprotonated uracil were generated following the antechamber protocol - as described in Supplementary Materials. The system was then soaked into a TIP3P^42^ water box with a 10 Å buffer and potassium ions to ensure a neutral global charge, resulting in a total of ∼ 181,000 atoms. The ff14SB force field including the OL3 corrections^43^ for RNA were used to describe the protein and the nucleic acid. RNAP presents a zinc fingers which has been modeled using non-bonded parameters.^44^ The protonation states of the protein residues were assigned with respect to the pKa prediction made with propka3.1.^45^ Of note, a cysteine near to the active site (C622) was considered as deprotonated for it is involved in the coordination of the magnesium cluster.

Hydrogen Mass Repartitioning (HMR)^46^ was consistently applied to all hydrogens excluding water, thus allowing, in combination with Rattle and Shake, ^**?**^ the use of a 4 fs time-step to numerically solve the Newton equations of motion. Prior to the simulation, the system geometry was optimized in four 10,000 steps with gradually decreasing restraints on the backbone atoms of the protein and RNA. Each of these steps was followed by a short 1.2 ns equilibration with the same restraints in order to ensure a proper relaxation of the system. A 150 ns production run was then performed at 300K in the isotherm and isobaric (NTP) ensemble. The temperature was kept constant using the Langevin thermostat with a 1 ps^−1^ collision frequency, and the electrostatic interaction were treated using the Particle Mesh Ewald (PME)^47^ scheme with a 9 Å cutoff. A 10 kcal/mol constraint was imposed on the distance between the magnesium ions forming the cluster to compensate for the electrostatic repulsion and maintain the bimetallic active site in a pre-reactive conformation.

Structural analysis was performed using the cpptraj module of AMBER and the Curves+ program.^48^ A clustering of the trajectory was performed to extract the most abundant conformation sampled as starting structure for the subsequent QM/MM-MD simulations.

### Equilibrium QM/MM-MD simulation

The representative frame of the most populated cluster issued from classical MD simulations was taken as a starting point for the QM/MM approach. The QM partition was defined to encompass the triphosphate moiety and C5’ atom of the GTP, the C2’, C4’ and C3’ the 3’ uracil (rU9), the magnesium ions, and the K551, K798, D760, D761, D618, C622 residues - see Figure S1 for a representation of the QM partition. All the involved amino acids were truncated at the C*α* position, resulting in a total of 81 atoms described at the QM level with a total charge of -3 and in a singlet spin state. The rest of the system was described at the MM level. Dangling covalent bonds have been treated with the link atom (hydrogen) approach.^**?**^ QM calculations were performed at Density Functional Theory (DFT) level using the *ω*B97x-D exchange and correlation functional^49^ and the double-*ζ* 6-31G basis set.^50^ The system was first relaxed at the QM/MM level for 10 ps, allowing the proper rearrangement of the active site and providing a relevant starting structure for the subsequent enhanced sampling free-energy calculation. The time-step for all the QM/MM simulations was set to 1 fs and all the Rattle and Shake constraints on the QM hydrogens were relaxed.

### Enhanced Sampling and Free Energy calculation

To allow for the exploration of chemical reactivity we resorted to enhanced sampling and more specific umbrella sampling (US) protocol as implemented in AMBER, thus obtaining a relevant free energy profile. Specifically, two collective variables were used to map the Free Energy Surface (FES) corresponding to the incorporation of GTP into the nascent RNA strand - see Figure 2. The first one (*ξ*) is defined as the difference between the distances U3’:O3’-GTP:PA (i.e. the phosphate forming bond) and GTP:PA-GTP:OP3 (i.e. the breaking bond). It is clear that *ξ* is a well adapted degree of freedom to describe the advancement of the nucleophilic substitution leading to the incorporation of the nucleotide in the RNA strand. As previously said proton shuttles are usually fundamental to assure polymerase efficiency, in particular assuring proton transfer to the pyrophosphate leaving group. To account for this, the second collective variable (*χ*) was defined as the difference between the distances K551:HZ1-GTP:O1B (forming bond) and K551:HZ1-K551:NZ (breaking bond), hence describing proton transfer from K551. US was performed partitioning the collective variables space into a regular grid spanning values of *ξ* from -2.0 to 1.8 Å and values of *χ* from -0.5 to 1.0 Å with steps of 0.1 Å. This resulted in 624 windows. For each window after a 1 ps equilibration 5 ps of production have been performed, for a total sampling of 3.2 ns. Values of *ξ* and *χ* were constrained on each window by harmonic potentials with anchors of strength of 600 kcal/mol set at -10 and 10 Å. The results were unbiased and FES reconstructed using the 2D version of the weighted histogram analysis method (WHAM2D).^51^ The consistence of the WHAM2D procedure was confirmed by checking the proper overlap between the windows as shown in Figure 2.

**Figure 2:**
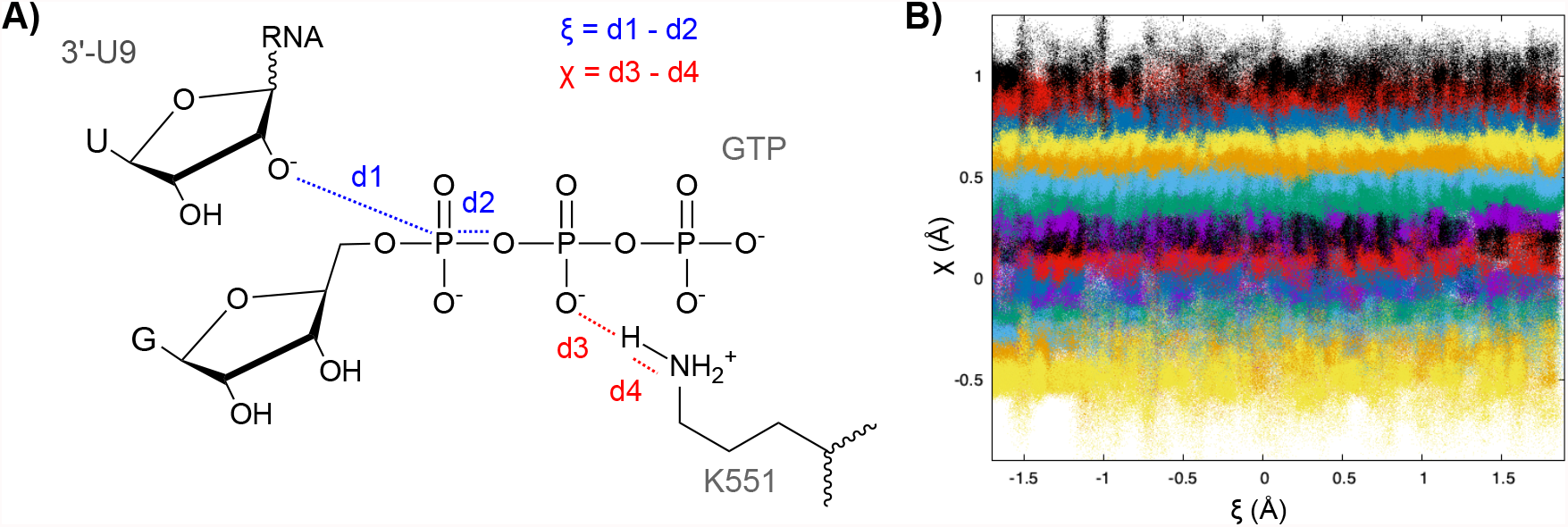
A) schematic definition of the collective variables *ξ* and *χ* used for the US simulation. B) Distribution of the collective variable values for all the simulated windows showing the overlap between them.

## Results and discussion

### Structural features of SARS-CoV-2 RNAP

The structural dynamic behavior of the RNAP:RNA complex was sampled by all-atom classical MD simulations. In particular in the following we describe the interaction network, leading to a stable protein/nucleic acid complex, as well as the main structural deformations experienced by the RNA strand and the enzymatic site. Importantly, to assure the accuracy of the viral replication RNAP relies on the Watson and Crick coupling between a nascent RNA strand and its original template. Hence, the nucleic acid needs to assume a double-stranded secondary structure, while its precise tertiary folding is largely dictated by the interaction with the RNAP interface.

At the interface between the nucleic acid and the enzyme, basic and polar amino acids develop non-specific contact with the RNA backbone leading to a dense interaction network. The rather unspecific binding of RNA and DNA through electrostatic interactions with the negative backbone is common among different nucleic acid processing or compacting proteins, including endonucleases or histone-like agents.

On the RNAP template strand side, positively-charged (R569, R914, K577) and polar residues (Y689, S501, Q541, N497, N507) interact with the negatively-charged phosphate groups of the RNA backbone - see Figures 3 and S2. While only one specific interaction is found between the protein and a RNA sugar ring, involving D684 and rA13, the latter is highly conserved along the simulation. Interestingly, S682 makes contacts with the nucleobase facing the incorporating nucleotide (rC12) facilitating the Watson and Crick pairing. This evidence suggests, as reported in previous works,^24,35^ that this amino acid might play a role in the relative positioning of the template strand and the reacting nucleotide in the active site, hence being crucial for efficiency and accuracy of the replication.

**Figure 3:**
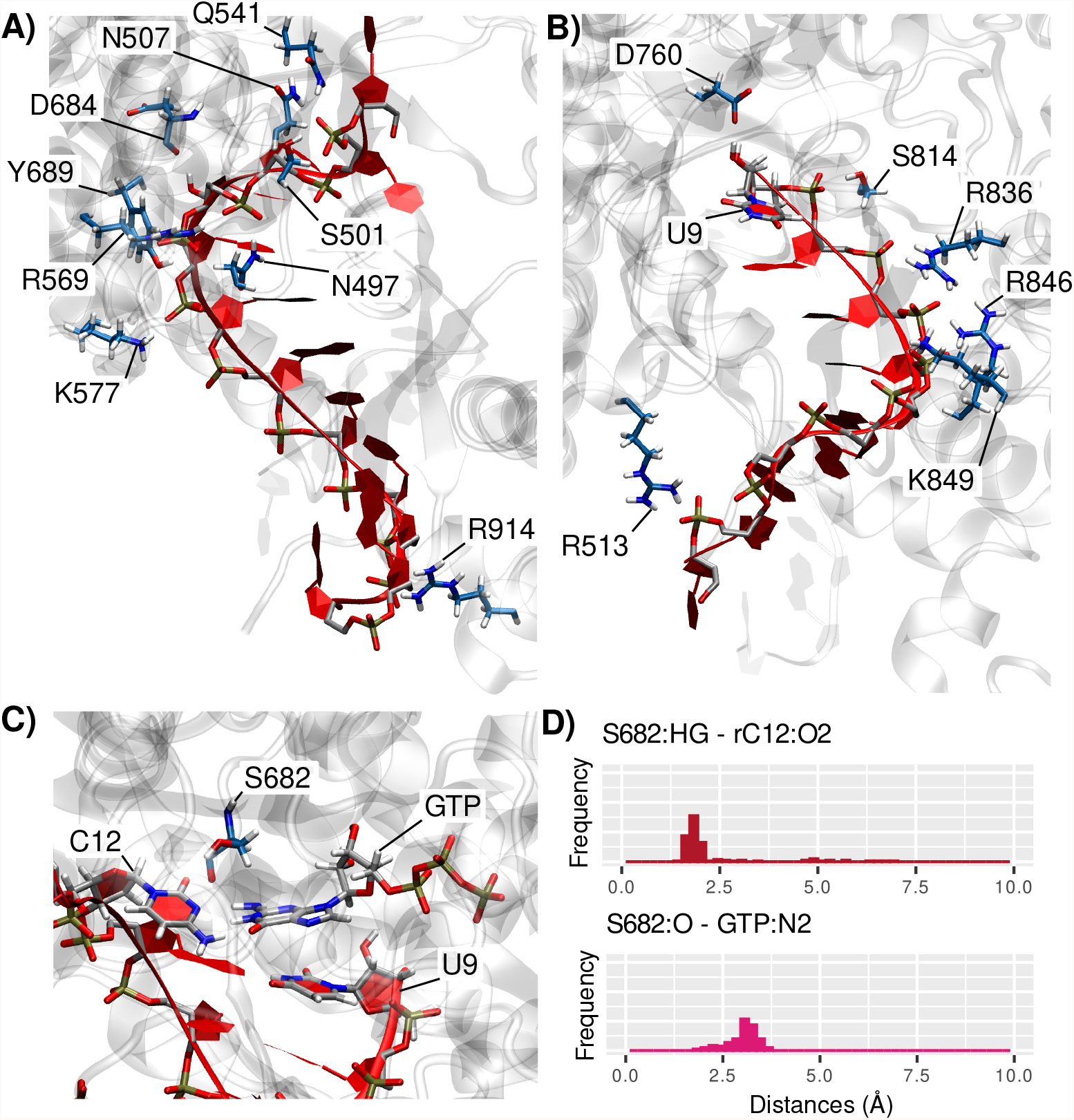
Main interactions between the enzyme (residues with blue carbon atoms) and A) the RNA template strand or B) the RNA nascent strand. The RNA strands are depicted in red and the protein structure appears in light grey. C) Zoom on the interaction of the key-residue S682 with the nucleotide to be integrated (GTP) and the facing rC12 on the template strand. D) Distribution of the distances between S682 and the base pair that is about to be formed.

Fewer interactions are observed involving the nascent strand. R513, R836, R858 and K849 are developing persistent contacts with the RNA phosphates, probably helping the correct positioning of the strand and moire importantly its translocation. Important interactions with the sugar ring of the terminal 3’-rU9 nucleotide are also observed that involve S814 and D760 - averaged distance of 2.25 ± 0.07 Å and 1.94 ± 0.04 Å, respectively. Interestingly, since the latter residue is part of the active site it is also coordinated to one of the catalytic magnesium ion.

Interestingly, the interaction between the double-strand nucleic acid and the enzyme, leading to a contact region spanning from base pair 5 to 9, is also translated into a rigidification and stabilization of the RNA. Indeed, the structural parameters of the RNA base-pairs as computed with Curves+ indicate a very stable and mostly ideal conformation of the double strand when in contact with the polymerase interface. This observation is most probably due to highly dense interaction network involving strong electrostatic interactions as above-described. Concerning the global structural descriptors of duplex RNA, the bending remains low for the base-pairs buried in the protein (up to 3.6 ± 2.2° for bp 5), while the regions more exposed to the solvent experience a higher flexibility with the local bending reaching 6.8 ± 6.7° for bp 2 - see Table 1. This behavior is also reflected in the systematically higher deviations of the axis parameters for the regions of the double strand which are more exposed to the water bulk - see Figure S3. Overall, the protein embedding stabilizes and rigidifies the RNA duplex structure, hence assuring the Watson and Crick coupling and offering an appropriate organization of the active site for the efficient incorporation of the reacting nucleotides into the nascent strand.

**Table 1:**
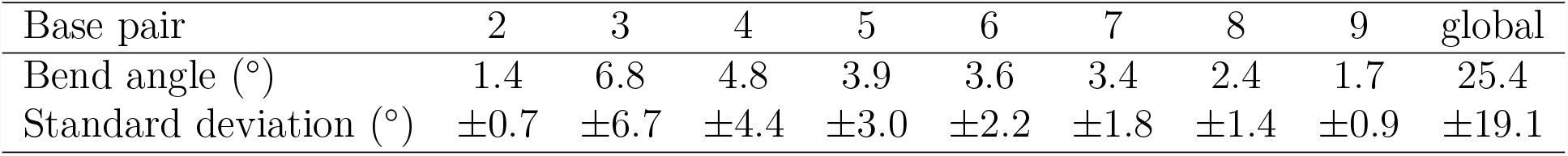
Averages and standard deviations of the local and global bend angles calculated along the classical MD trajectory.

As concerns the local organization of the active site, a first magnesium ion coordinates to D760, D618, the GTP reactive phosphate group (OP1), and the terminal deprotonated O3’ of the nascent RNA strand. This interaction clearly helps in maintaining the reactive moieties ideally placed for the reaction to take place - see Figure 1-A. The second Mg^2+^ ion interacts with the two other phosphates of GTP, D618, and D761, as well as the 3’ uracil phosphate group and O3’ atom through a bridging water. The sugar moiety of GTP is also involved in a hydrogen bond with C622. This highly organized electrostatic network within the active site, maintained by the presence of the 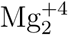 cluster, provides a nearly ideal structure favoring the nucleophilic substitution. As a matter of fact, the reactive rU9:O3’ and GTP:PA are kept only 3.5 Å apart, while the corresponding nucleobases develop an ideal *π*-stacking further stabilizing their arrangement. Notably, a canonical Watson-Crick hydrogen bonding scheme is observed between GTP and the matching rC12 on the template strand.

As already pointed out the active site of RNAP lacks a histidine, which could potentially participate in the proton transfer necessary to the completion of the reaction. While the analysis of the crystal structure of RNAP does not allow to identify any suitable amino acid nearby the 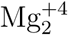 capable of assuming this role, the evolution of the MD simulation reveals that a proton donor, namely K551, is rapidly approaching the pyrophosphate leaving group, and is then stabilized in this conformation. K551 is not directed towards the active site in the crystal structure of RNAP with Favipiravir, however when the ligand is mutated to GTP a reorganization takes place in which K551 forms strong hydrogen bonds with the pyrophosphate group (K551:HZ2-GTP:O2G at 1.74 Å). Therefore we may hypothesize that K551 may compensate for the absence of the histidine, which, in turn, suggest an important role of this residue in SARS-CoV-2 RNA polymerase enzymatic mechanism. This hypothesis will be further confirmed also by our QM/MM simulations. Of note, this lysine is highly conserved in other viral polymerase, hence suggesting a conserved biological role of this residue - see Figure S4. Importantly the main characteristic of the active site of RNAP in complex with GTP are also retained during the QM/MM equilibration, in which only slightly adaptation of the Mg-Mg distance can be pointed out.

### Reaction Free Energy Profile

The free energy profile of the enzymatic reaction catalyzed by RNAP was obtained to precisely quantify the associated activation energy and driving force. The common reaction mechanism for polymerases is based on a two-steps process: i) the activation of the 3’ nucleotide of the nascent strand through its deprotonation, also coupled with the protonation of the leaving pyrophosphate group - see Figure 1-C; ii) the nucleophilic attack involving the activated 3’-nucleotide and the incorporating triphosphate nucleotide, resulting in the elongation of the nascent RNA strand and the release of the pyrophosphate group. Here, only the rate limiting step of the global reaction i.e., the nucleophilic attack, was simulated by QM/MM-MD calculations. The free energy surface was determined along two reaction coordinates simultaneously describing the attack of rU9:O3’ onto the GTP phosphate (*ξ*), and the assistance and potential proton transfer of the -NH3^+^ moiety of K551 facilitating the pyrophosphate departure (*χ*). Indeed, K551 has been pinpointed by structural consideration and sequence alignment as a key residue for the polymerase reactivity. Equilibrium MD simulations on the protonated U9:O3’ show a high water accessibility of the OH group, with at least one solvent molecule present in the closest coordination shell of rU9:O3’ for 72% of the global simulation time. The solvent exposition and the usual lability of alcohol moieties contribute to suggest a facile deprotonation of the 3’-terminal nucleotide, and support our choice to concentrate on the second step only.

The 2D free energy surface along *ξ* and *χ* is reported in Figure 4-A, together with representative snapshots of critical points of the surface (4-B). The reaction exhibits a classical nucleophilic substitution mechanism, going from the reactant (R) to the product (P) through a transition state (TS) without any intermediate. The TS is accessed bypassing only a moderate to low activation barrier ΔG^≠^ of 10 kcal/mol, hence confirm the efficiency of RNAP. Additionally, the TS exhibits a perfectly ideal planar reacting phosphate group, which is also equidistant from O3’ and O3A (1.9 Å), as can be appreciated from the evolution of the distance and the phosphate pyramidalisation angle reported in Figure 4-B and C. From a thermodynamic point of view, the reaction is found to be exergonic, with a driving force of ΔG^0^=18 kcal/mol. Upon reaching the TS, the distance between the catalytic Mg^2+^ ions is slightly increased from 4.2 Å to 4.8 Å. The first ion maintains its interaction with D760 and D618, but not anymore with the rU9:O3’ which is no longer accessible and is instead coordinated to the phosphate’s oxygen of the newly incorporated guanine monophosphate. The second ion also interacts with the the backbone of the new 3’ nucleotide, D168, and one end of the pyrophosphate.

**Figure 4:**
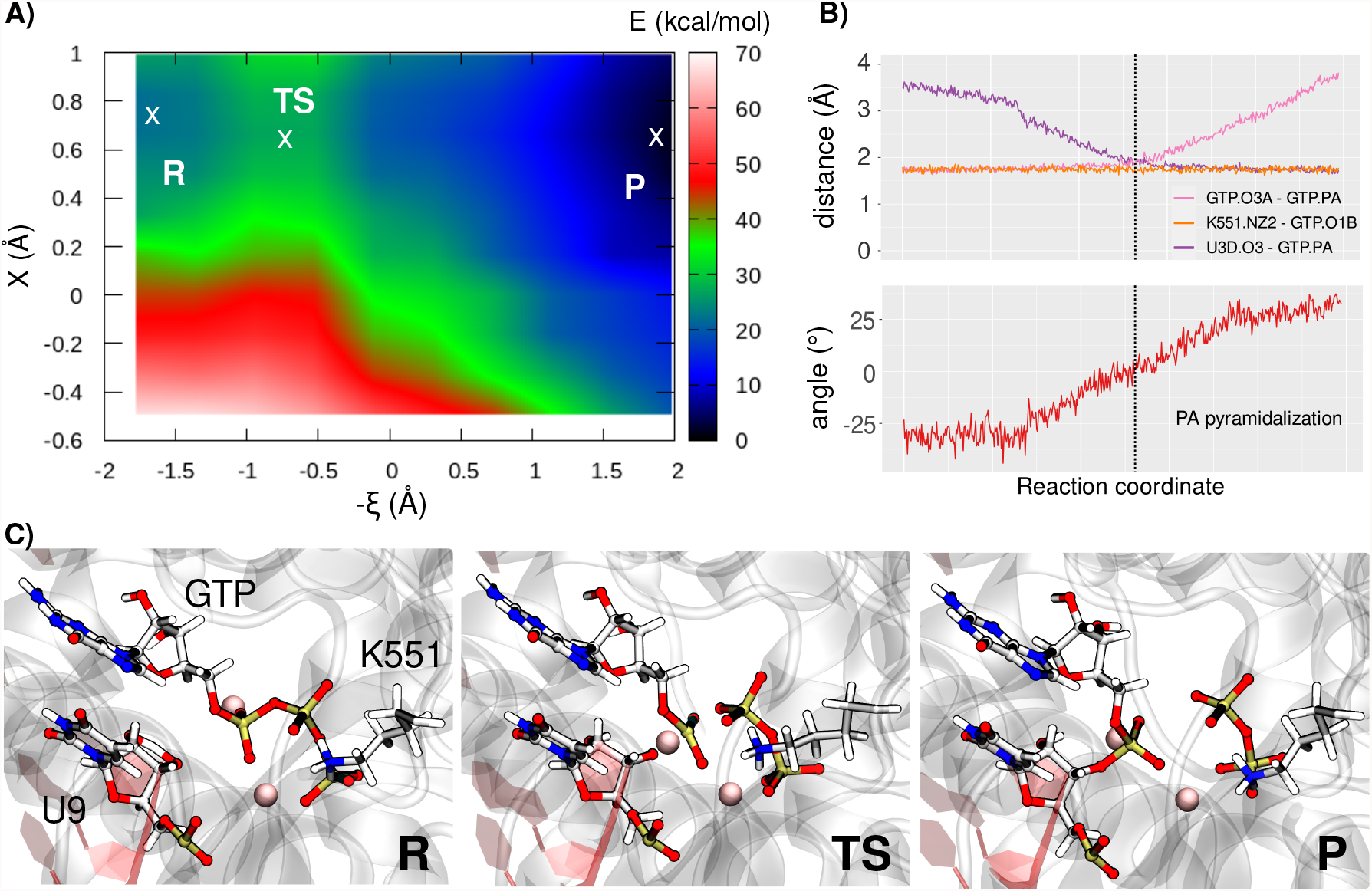
A) 2D Free energy surface obtained from the QM/MM umbrella sampling simulations. B) Evolution along the reaction coordinate *ξ* of the distances between GTP and protein or RNA moieties (upper panel) and of the pyramidalisation angle of the reacting phosphate (lower panel). Note that due to the topology of the free energy surface only value of *χ* = 0.7 Å have been considered. C) Representative snapshots describing the critical points observed on the free energy maps, the position of the reactants (R), transition state (TS), and products (P) in the free energy basins is also provided in panel A

Interestingly, while the proton transfer from K551 to the pyrophosphate does not appear to be required for the completion of the reaction, as can be appreciated by the topology of the free energy surface, and by the fact that the minimum energy path lays around values of *χ*=0.7 Å. However, the role of K551 is crucial as it induces an important stabilization of the reaction product. Interestingly, the lysine proton is much more labile in the product region, and, in shark contrast with the reactants, the presence of a large low-energy basin spanning values of *χ* comprised between -0.6 and 1.0 Å should be highlighted. Hence, once the nucleotide is incorporated into the nascent strand, K551 is able to partially share its proton with the leaving pyrophosphate group facilitating its departure. This peculiar topology of the free energy surface, also confirms the different nature of RNAP enzymatic reactions. Indeed, the absence of the histidine precludes the possibility of a proton-shuttle mechanism, however a lysine, in this case K551, may act as a partial proton donor leading to a product region in which the additional charge is largely delocalized over different moieties. To the best of our knowledge, this peculiar mechanism has not been identified in other polymerases, and shows how unusual active sites may lead to highly efficient turnovers.

Starting from the structure of the reaction product (window *ξ*=2.0 and *χ*=0.7), an additional unbiased QM/MM-MD trajectory of 100 ps was performed to sample at least the initial phases of the pyrophosphate group departure. In the course of the unbiased dynamics K551 continues to interact with the leaving group and interestingly spontaneously drags it towards the bulk. Besides K551, other basic amino acids, namely K798, R553, and R555, approach the active site and develops interactions with the pyrophosphate globally favoring its departure from the active site - see Figure 5-A.

**Figure 5:**
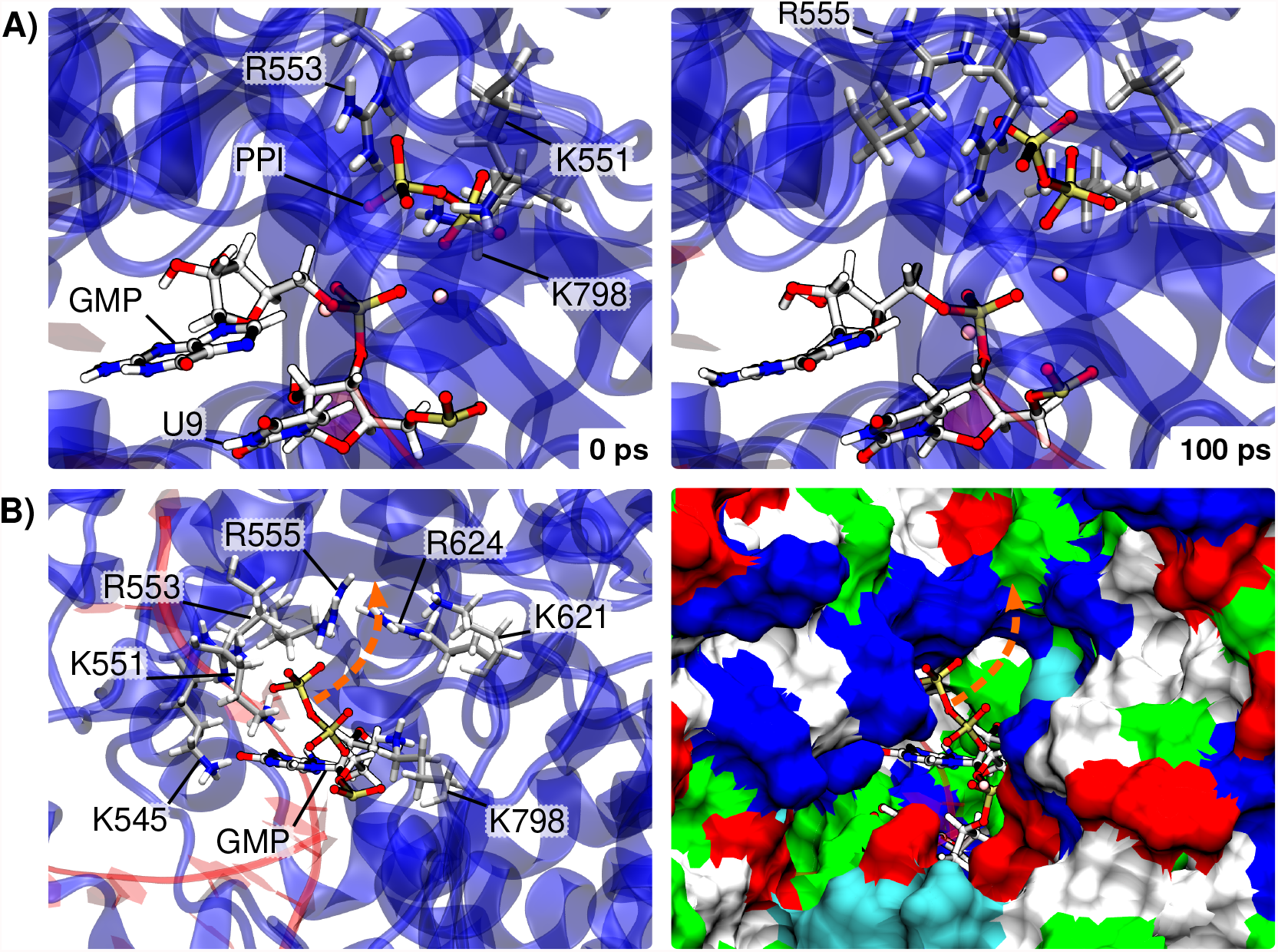
A) Structure of RNAP active site at the beginning (0 ps, left) and end (100 ps, right) of the unbiased QM/MM-MD simulations of RNAP after GMP incorporation into the nascent strand. The pyrophosphate (PPI) and basic amino acids within 5 Å are displayed, showing the movement of PPI upwards, moving closer to R555 towards the exit. B) Global view of the basic amino acids within 10 Å of the PPI moiety *en route* towards the exit (after 100 ps unbiased sampling). On the left the basic amino acids are explicitly depicted in licorice, and on the right a surface representation of the protein is used with a color code of the amino acids corresponding to their type (basic in blue) highlighting the the possible exit path for PPI. This path appears as an orange arrow on both representations.

Since in our simulation, one of the phosphates remains coordinated to a magnesium ion, we have not observed a complete detachment of the leaving group. However, we may anticipate that longer simulations might show a facile complete exit of the pyrophosphate from the active site, also thanks to the development of the interaction networks with the basic amino acids. The inspection of the basic amino acids and surface of the protein offering a possible exit path for PPI show an ideal way upwards, with many lysines and arginines (K551, R553, R555, R624, K798) that might favor the passage of the leaving group towards the bulk - see Figure 5-B. Interestingly, Naydenova et al. report a nonproductive conformation of RNAP harboring Favipiravir and a pyrophosphate above it that interacts with R555 and K621.^35^ They suggest that this pyrophosphate group might be a by-product from previous incorporation of NTPs into the nascent RNA strand. Its position corroborates the exit route that we hypothesize here.

## Conclusion

RNAP is a crucial enzyme in the viral cycle of SARS-CoV-2, and more generally RNA dependent RNA polymerases constitute a crucial target for the design of inhibitors against emerging viruses. While the structure of RNAP, in complex with different inhibitors, has been successfully resolved since the first outbreaks of the COVID-19 pandemics, its catalytic mechanism has not yet been fully elucidated from a biochemical point of view. With this contribution, following a multiscale approach combining classical and QM/MM simulations including enhanced sampling, we provide for the first time the description of the catalytic mechanism of RNAP, while quantifying the activation energy necessary for the RNA polymerization. From a structural point of view RNAP presents a high density of basic amino acids, producing a positively charged groove which is necessary to accommodate, in a rather unspecific way, the Watson and Crick-paired nascent and template strands. Interestingly, some specific interactions locking the template base in an ideal position to interact with the incorporated nucleotide are observed, thus increasing the specificity. If the RNAP active site presents many of the classical features of polymerases, including a 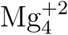 cluster, it lacks histidines susceptible to assist the polymerization by inducing a proton transfer to the leaving pyrophosphate group. The combination of classical and QM/MM-MD simulations has allowed to highlight the role of K551 in compensating for the absence of a histidine and in stabilizing the leaving pyrophosphate with the assistance of one of its protons. In particular we have evidenced that, while the polymerization reaction necessitates an activation energy of only 10 kcal/mol^−1^, the presence of K551 is also fundamental to favor the departure of the pyrophosphate, also assisted by other basic amino acids. Thus, we may conclude that RNAP is characterized by a high enzymatic efficiency, which could translate in a high viral replication rate, even if it presents an unusual active site compared to other similar proteins, and hence a partially different chemical mechanism. Our results, in providing a clear structural and mechanistic picture of the polymerization reaction, may open the way to the design of specific inhibitors, mainly based on nucleotide analogs. In particular, the key role played by K551 may also open possibilities for the design of inhibitors specifically targeting this residue. In the future we will extend this study to the reactivity of RNAP in presence of inhibitors, such as Remdesivir and Favipiravir, in order to assess the different modes of actions of these two compounds. Furthermore, the protocol presented here can be easily extended also to the study of other RNA-dependent RNA polymerases and hence can allow targeting other viral systems.

## Supporting information

Movie of the reaction mechanism

## Acknowledgement

The authors thank GENCI, Explor, and local LPCT computing centers for computational resources. E.B. thanks the CNRS and French Ministry of Higher Education Research and Innovation (MESRI) for her postdoc fellowship under the GAVO program. AM thanks ANR and CGI for their financial support of this work through Labex SEAM ANR 11 LABX 086, ANR 11 IDEX 05 02. The support of the IdEx “Université Paris 2019” ANR-18-IDEX-0001 and of the Platform P3MB is gratefully acknowledged.

## Supporting Information Available

The following information are available free of charge. Parameters and parameterization procedure used to define the non-standard nucleic bases used in this study. Representation of the atoms in the QM partition. Additional structural parameters for the RNA double strand obtained using Curves+. Movie extracted from the US simulation representing the structural evolution during the polymerase reaction.

## TOC Graphic

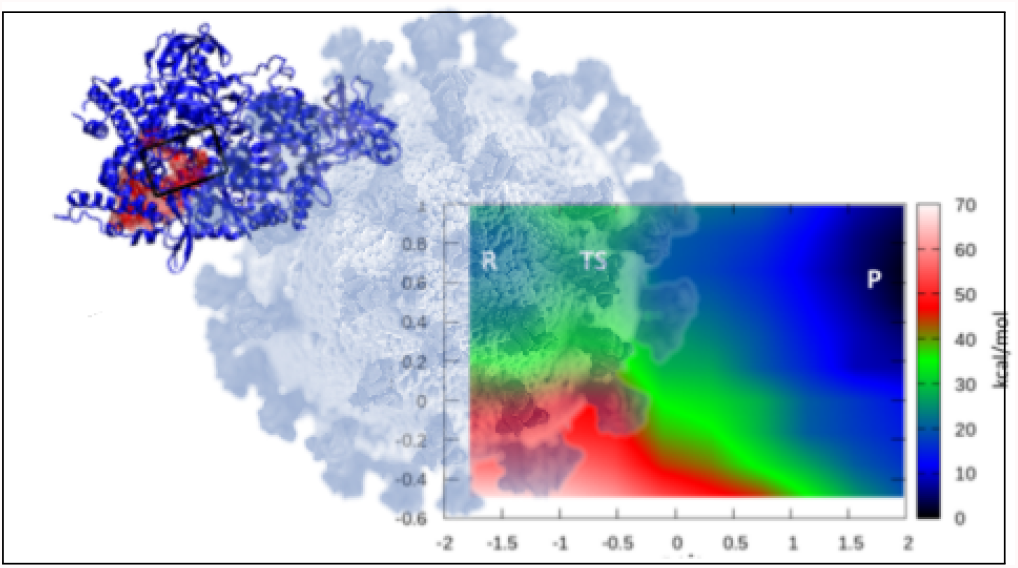

## Notes

### Competing Interest Statement

The authors have declared no competing interest.

