## Supplementary material for "Unraveling the Enzymatic Mechanism of the SARS-CoV-2 RNA-Dependent-RNA-Polymerase. An Unusual Active Site Leading to High Replication Rates": SI

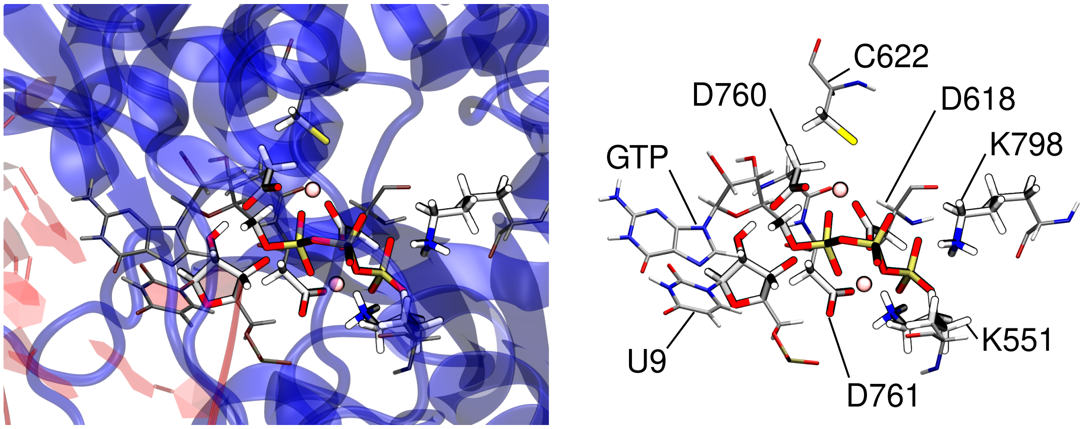


Figure S1 – Representations of the atoms involved in the QM partition of the QM/MM calculations. The QM atoms appear brighter and thicker, the rest of the residues' atoms are represented as thinner and darker to appreciate the boundary between the QM and MM partitions. On the left, the protein and nucleic embedding are depicted in blue and red, respectively, and on the right only the residues involved in the QM part are represented for sake of clarity. The magnesium ions appear as pink circles.

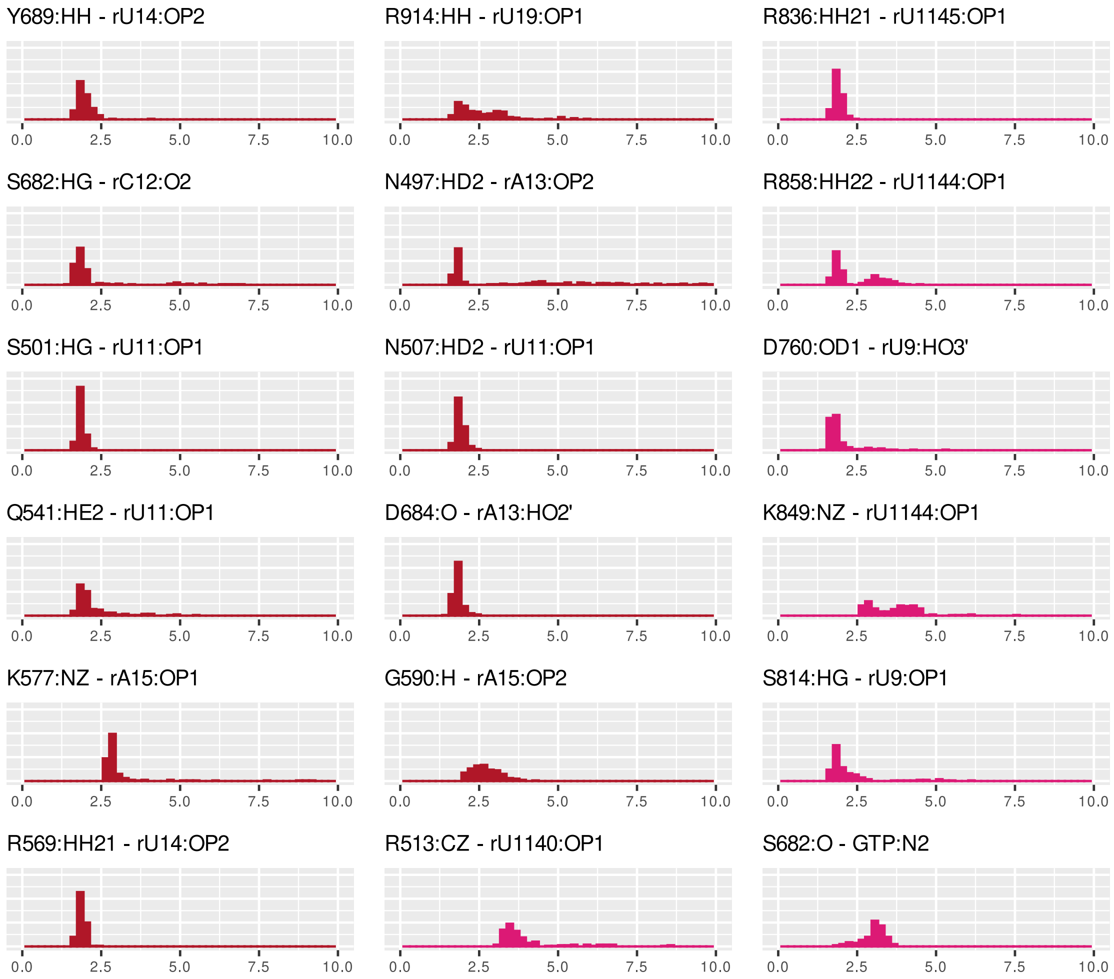


Figure S2 – Distribution of the distances characterizing the main interactions between the enzyme and the RNA duplex. Interactions with the template or nascent strands appear in red and pink, respectively.

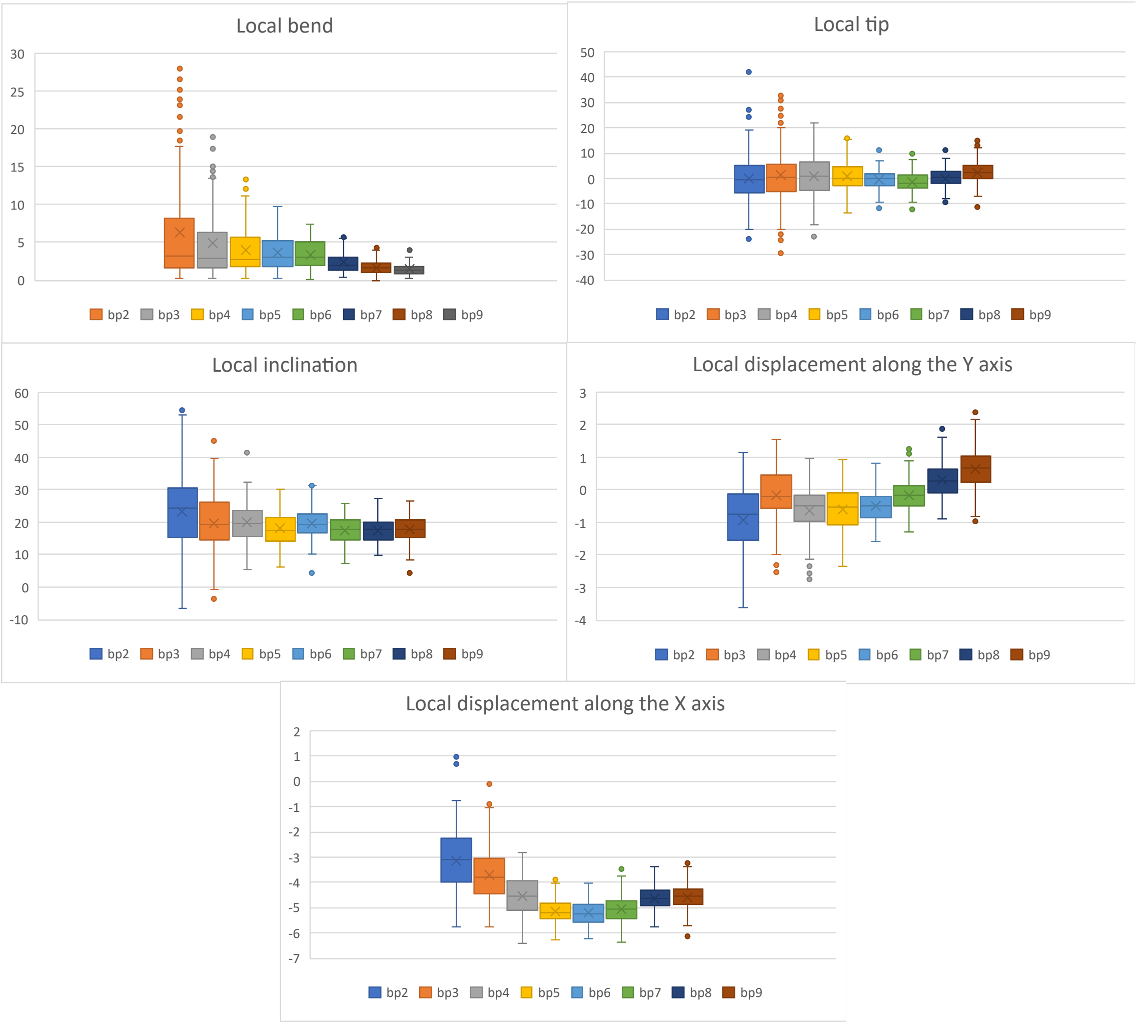


Figure S3 – Local axis parameters for base pairs 2 to 9, as computed by Curves+. Base pairs 2 to 4 are more exposed to the solvent while the pairs 5 to 9 are buried within the polymerase.

```
sp|P0DTD1|R1AB_SARS2      PFNKWGKARLYYDSMSYEDQDALFAYTKRNVIPTITQMNLKYAISAKNRARTVAGVSICS  4956
sp|P0C6X7|R1AB_SARS       PFNKWGKARLYYDSMSYEDQDALFAYTKRNVIPTITQMNLKYAISAKNRARTVAGVSICS  4933
sp|K9N7C7|R1AB_MERS1      PFNKFGKARVYYESMSYQEQDELFAMTKRNVIPTMTQMNLKYAISAKNRARTVAGVSILS  4942
sp|P0C6Y3|R1AB_IBVM       PFNKFGKARLYYE-MSLEEQDQLFESTKKNVLPTITQMNLKYAISAKNRARTVAGVSILS  4502
sp|P0C6Y5|R1AB_CVPPU      PLNKFGKARLYYETLSYEEQDALFALTKRNVLPTMTQMNLKYAISGKARARTVGGVSLLS  4559
sp|P0C6X1|R1AB_CVH22      PLNKFGKAGLYYESISYEEQDAIFSLTKRNILPTMTQLNLKYAISGKERARTVGGVSLLA  4627
sp|P0C6V9|R1AB_BC279      PFNKWGKARLYYDSMSYEDQDVLFAYTKRNVIPTITQMNLKYAISAKNRARTVAGVSICS  4939
sp|P0C6W2|R1AB_BCHK3      PFNKWGKARLYYDSMSYEDQDALFAYTKRNVIPTITQMNLKYAISAKNRARTVAGVSICS  4927
sp|P0C6W6|R1AB_BCRP3      PFNKWGKARLYYDSMSYEDQDALFAYTKRNVIPTITQMNLKYAISAKNRARTVAGVSICS  4931
                          *:**:*** :**: :* ::** :*  **:*::**:**:*******.* *****.***: :
```

Figure S4 – Multiple Sequence Alignment obtained with the Clustal Omega webserver, centered around the lysine corresponding to K551 in SARS-CoV-2 to highlight its conservation among other viruses' polymerase (in yellow). SARS2 = Severe acute respiratory syndrome coronavirus 2; SARS = Severe acute respiratory syndrome coronavirus; MERS1 = Middle East respiratory syndrome-related coronavirus; IBVM = Avian infectious bronchitis virus; CVPPU = Porcine transmissible gastroenteritis coronavirus; CVH22 = Human coronavirus 229E; BC279 = Bat coronavirus 279/2005; BCHK3 = Bat coronavirus HKU3; BCRP3 = Bat coronavirus Rp3/2004.
